# Traces of semantization - from episodic to semantic memory in a spiking cortical network model

**DOI:** 10.1101/2021.07.18.452769

**Authors:** Nikolaos Chrysanthidis, Florian Fiebig, Anders Lansner, Pawel Herman

## Abstract

Episodic memory is the recollection of past personal experiences associated with particular times and places. This kind of memory is commonly subject to loss of contextual information or “semantization”, which gradually decouples the encoded memory items from their associated contexts while transforming them into semantic or gist-like representations. Novel extensions to the classical Remember/Know behavioral paradigm attribute the loss of episodicity to multiple exposures of an item in different contexts. Despite recent advancements explaining semantization at a behavioral level, the underlying neural mechanisms remain poorly understood. In this study, we suggest and evaluate a novel hypothesis proposing that Bayesian-Hebbian synaptic plasticity mechanisms might cause semantization of episodic memory. We implement a cortical spiking neural network model with a Bayesian-Hebbian learning rule called Bayesian Confidence Propagation Neural Network (BCPNN), which captures the semantization phenomenon and offers a mechanistic explanation for it. Encoding items across multiple contexts leads to item-context decoupling akin to semantization. We compare BCPNN plasticity with the more commonly used spike-timing dependent plasticity (STDP) learning rule in the same episodic memory task. Unlike BCPNN, STDP does not explain the decontextualization process. We also examine how selective plasticity modulation of isolated salient events may enhance preferential retention and resistance to semantization. Our model reproduces important features of episodicity on behavioral timescales under various biological constraints whilst also offering a novel neural and synaptic explanation for semantization, thereby casting new light on the interplay between episodic and semantic memory processes.

## 1 INTRODUCTION

Remembering single episodes is a fundamental attribute of human cognition. A memory, such as with whom you celebrated your last birthday, is more vividly recreated when we can recall contextual information, such as the location of the event (Eichenbaum et al., 2007; Gillund, 2012). The term “episodic memory” was originally introduced by Tulving (1972) to designate memories of personal experiences. Retrieval from episodic memory includes a feeling of mental time travel realized by “I remember”. In contrast, semantic memory retrieval encapsulates what is best described by “I know” (Tulving, 1985; Umanath and Coane, 2020). Unlike episodic memories, semantic memories refer to general knowledge about words, items and concepts, lacking spatiotemporal source information, possibly resulting from the accumulation of episodic memories (Schendan, 2012; Gillund, 2012).

Initially, Tulving (1972) proposed that episodic and semantic memory are distinct systems and compete in retrieval. Recent studies suggest, however, that the division between episodic and semantic memory is rather vague (McCloskey and Santee, 1981; Renoult et al., 2019), as neural activity reveals interaction between episodic and semantic systems during retrieval (Weidemann et al., 2019). According to Squire and Zola (1998) retrieval of semantic memory depends on the acquisition of the episode in which such information was experienced. Apparently, there is a clear interdependence between the two systems as the content of episodic memory invariably involves semantic representations (Martin-Ordas et al., 2014), and consequently semantic similarity aids episodic retrieval (Howard and Kahana, 2002).

Episodic memory traces are susceptible to transformation and loss of information (Tulving, 1972), and this loss of episodicity can be attributed to semantization, which typically takes the form of a decontextualization process (Duff et al., 2020; Habermas et al., 2013; Viard et al., 2007). Meeter and Murre (2004) highlight and review the dynamical nature of memories and neural interactions through the scope of Transformation theory, which suggests that all memories start as episodic representations that gradually transform into semantic or gist-like representations (Winocur and Moscovitch, 2011; Petrican et al., 2010). Decontextualization can occur over time as studies suggest that older adults report fewer episodic elements than younger adults (Petrican et al., 2010). Yet, could this item-context decoupling rely on accumulation of episodicity over multiple exposures of stimuli in various contexts over time? Baddeley (1988) hypothesized that semantic memory might represent the accumulated residue of multiple learning episodes, consisting of information which has been semanticized and detached from the associated episodic contextual detail. In fact, simple language vocabulary learning implies that learners encode words in several different contexts, which leads to semantization and definition-like knowledge of the studied word (Beheydt, 1987; Bolger et al., 2008).

Retrieval from episodic memory has been studied extensively through the lens of the classical Remember/Know (R/K) paradigm, in which participants are required to provide a Know or Remember response to stimulus-cues, judging whether they are able to recall item-only information or additional details about episodic context, respectively (van den Bos et al., 2020). Extensions of the classical R/K behavioral experiment demonstrate that item-context decoupling can occur rapidly (Opitz, 2010). In these experiments, items are presented during an encoding phase either in a unique context, or across several contexts. Low context variability leads to greater recollection, whereas context overload results in decontextualization and a higher fraction of correctly classified Know responses (Opitz, 2010; Smith and Manzano, 2010; Smith and Handy, 2014). In the current study, we offer and evaluate a Bayesian-hypothesis about synaptic and network mechanisms underlying the memory semantization (item-context decoupling).

In earlier works, we developed and investigated a modular spiking neural network model of cortical associative memory with respect to memory recall, including oscillatory dynamics in multiple frequency bands, and compared it to experimental data (Lundqvist et al., 2010, 2011; Herman et al., 2013). Recently we demonstrated that the same model, enhanced with a Bayesian-Hebbian learning rule (Bayesian Confidence Propagation Neural Network, BCPNN) to model synaptic and intrinsic plasticity, was able to quantitatively reproduce key behavioral observations from human word-list learning experiments (Fiebig and Lansner, 2017), such as serial order effects during immediate recall. This model performed one-shot memory encoding and was further expanded into a two-area cortical model used to explore a novel indexing theory of working memory, based on fast Hebbian synaptic plasticity (Fiebig et al., 2020). In this context, it was suggested that the underlying naive Bayes view of association would make the associative binding between two items weaker if one of them is later associated with additional items. Thus, if we conceive of episodicity as an associative binding between item and context, the BCPNN synaptic plasticity update rule might provide a mechanism for semantization. In this work, we test this hypothesis and examine to what extent the results match available behavioral data on semantization. We further compare those outcomes of dynamic learning with a model featuring the more well-known spike-timing dependent plasticity (STDP) learning rule. We also demonstrate how selective plasticity modulations of one-shot learning (tentatively modelling effects of attention, emotional salience, valence, surprise, etc. on plasticity) may enhance episodicity and counteract semantization.

To our knowledge, there are no previous computational models of item-context decoupling akin to semantization. Overall, there are rather few computational models of episodic memory (Norman and O’Reilly, 2003), and those that exist are typically abstract, aimed at predicting behavioral outcomes without a specific focus on underlying neural and synaptic mechanisms (Greve et al., 2010; Wixted, 2007). Our model bridges these perspectives and explains semantization based on synaptic plasticity, while it also reproduces important episodic memory phenomena on behavioral time scales under constrained network connectivity with plausible postsynaptic potentials, firing rates, and other biological parameters.

## 2 RESULTS

### 2.1 Semantization of episodic representations in the BCPNN model

The network model used here features two reciprocally connected networks, the so-called Item and Context networks. The architecture of each network follows our previous spiking implementations of attractor memory networks (Lansner, 2009; Tully et al., 2014, 2016; Lundqvist et al., 2011; Fiebig and Lansner, 2017; Chrysanthidis et al., 2019; Fiebig et al., 2020), and is best understood as a subsampled cortical layer 2/3 patch with nested hypercolumns (HCs) and minicolumns (MCs; Fig. 1A, see STAR⋆METHODS for details). In our model, items are embedded in the Item network, and context information in the Context network as internal long-term memory representations, derived from prior Hebbian learning (Fig. 1B,C, STAR⋆METHODS). Our episodic memory task is designed to follow a seminal experimental study by Opitz (2010). We stimulate some items in a single context and others in a few different contexts establishing multiple associations (Fig. 2). Stimulus duration during encoding is *t*_*stim*_=250 ms with a *T*_*stim*_=500 ms interstimulus interval, and a test phase occurs after a 1 s delay period, which contains brief *t*_*cue*_=50 ms cues of previously learned items (Table S2).

**Fig. 1.**
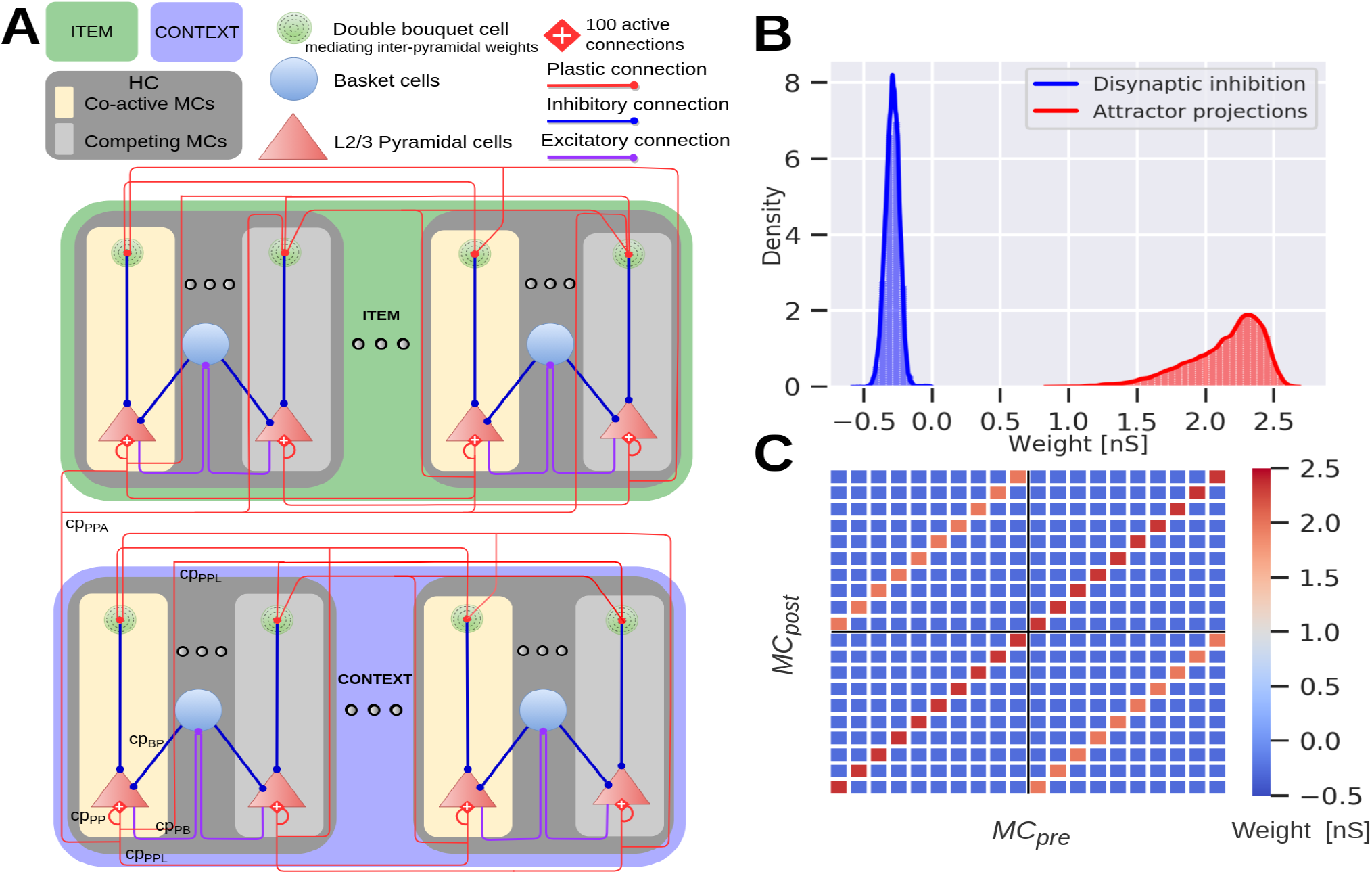
Network architecture and connectivity of the Item (green) and Context (blue) networks. **A)** The model represents a subsampled modular cortical layer 2/3 patch consisting of minicolumns (MCs) nested in hypercolumns (HCs). Both networks contain 12 HCs, each comprising 10 MCs. We preload abstract long-term memories of item and context representations into the respective network, in the form of distributed cell assemblies with weights establishing corresponding attractors. Associative plastic connections bind items with contexts. The network features lateral inhibition via basket cells (purple and blue lines) resulting in a soft winner-take-all dynamics. Competition between attractor memories arises from this local feedback inhibition together with disynaptic inhibition between HCs. **B)** Weight distribution of plastic synapses targeting pyramidal cells. The attractor projection distribution is positive with a mean of 2.1, and the disynaptic inhibition is negative with a mean of -0.3 (we show the fast AMPA weight components here, but the simulation also includes slower NMDA weight components). **C)** Weight matrix between attractors and competing MCs across two sampled HCs. The matrix displays the mean of the weight distribution between a presynaptic (*MC*_*pre*_) and postsynaptic minicolumn (*MC*_*post*_), within the same or different HC (black cross separates grid into blocks of HCs, only two of which are shown here). Recurrent attractor connections within the same HC are stronger (main diagonal, dark red) compared to attractor connections between HCs (off-diagonals, orange) while inhibition is overall balanced between patterns (blue). Negative inter-pyramidal weights between competing MCs amounts to disynaptic inhibition mediated by double bouquet cells.

**Fig. 2.**
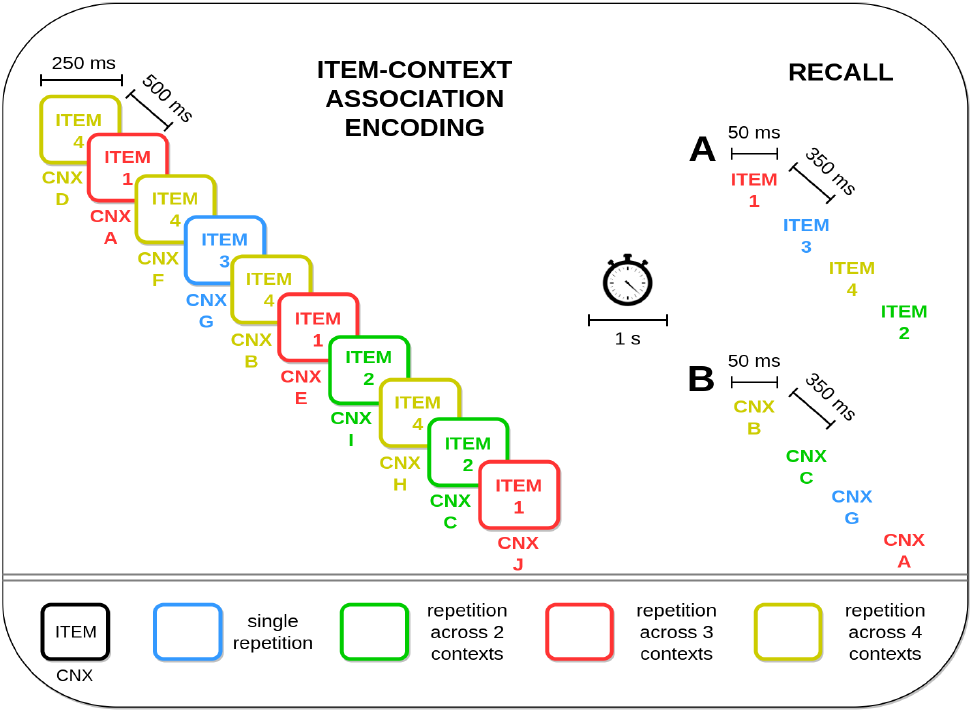
Trial structure of the two simulated variants of the episodic memory task. Items are first associated with one or several contexts (CNX) during the encoding phase in 250 ms cue episodes, with an interstimulus interval of 500 ms. The colors of the co-activated contexts are consistent with their corresponding associated item. The recall phase occurs with a delay of 1 s and involves different trials with either brief cues (50 ms) of the **A)** items, or **B)** contexts presented during the item-context association encoding phase.

Figure 3A illustrates an item-context pair, established by an associative binding through plastic bidirectional BCPNN projections (dashed lines). Item and context attractors (solid red lines) are embedded in each network and remain fixed throughout the simulation, representing well-consolidated long-term memory. We show an exemplary spike raster of pyramidal neurons in HC1 of both the Item and Context networks reflecting a trial simulation (Fig. 3B). Herein, item-3 (blue) establishes a single association, whilst item-4 (yellow) is encoded in four different contexts (Fig. 2A, 3B). We observe evidence of item-context decoupling as the yellow item (but not the blue) is successfully recognized when cued but without any corresponding accompanying activation in the Context network. Successful and complete item recognition without any contextual information retrieval accounts for a Know response, as opposed to Remember judgments, which are accompanied by successful context recall. Cue-based activations are reported using a detection algorithm (see STAR⋆METHODS). Figure 3C demonstrates the performance of contextual retrieval when items serve as cues. To elucidate this observed progressive loss of episodicity, we sample and analyze the learned weight distributions of item-context binding recorded after the association encoding period (Fig. 3D). The item-context weight distribution in the one-association case has a significantly higher mean than in the two-, three-, or four-association case (*p*<0.001, Mann-Whitney, *N*=2000). This progressive weakening of weights leads to significantly lower mean EPSP amplitudes for the associative projections (*p*<0.05 for one vs two associations; *p*<0.001 for two vs three and three vs four associations, Mann-Whitney, *N*=300, Fig. S1). So, we attribute the loss of episodicity to a statistically significant weakening of means of the associative weight distributions with the increasing number of associated contexts. The associative weight distributions shown here refer to the NMDA component, while the weight distributions of the faster AMPA receptor connections display a similar trend (Fig. S2). The gradual trace modification we observe relies on the nature of Bayesian learning, which normalizes and updates weights over estimated presynaptic (Bayesian-prior) as well as postsynaptic (Bayesian-posterior) spiking activity (see Sect. 2.3 for details).

**Fig. 3.**
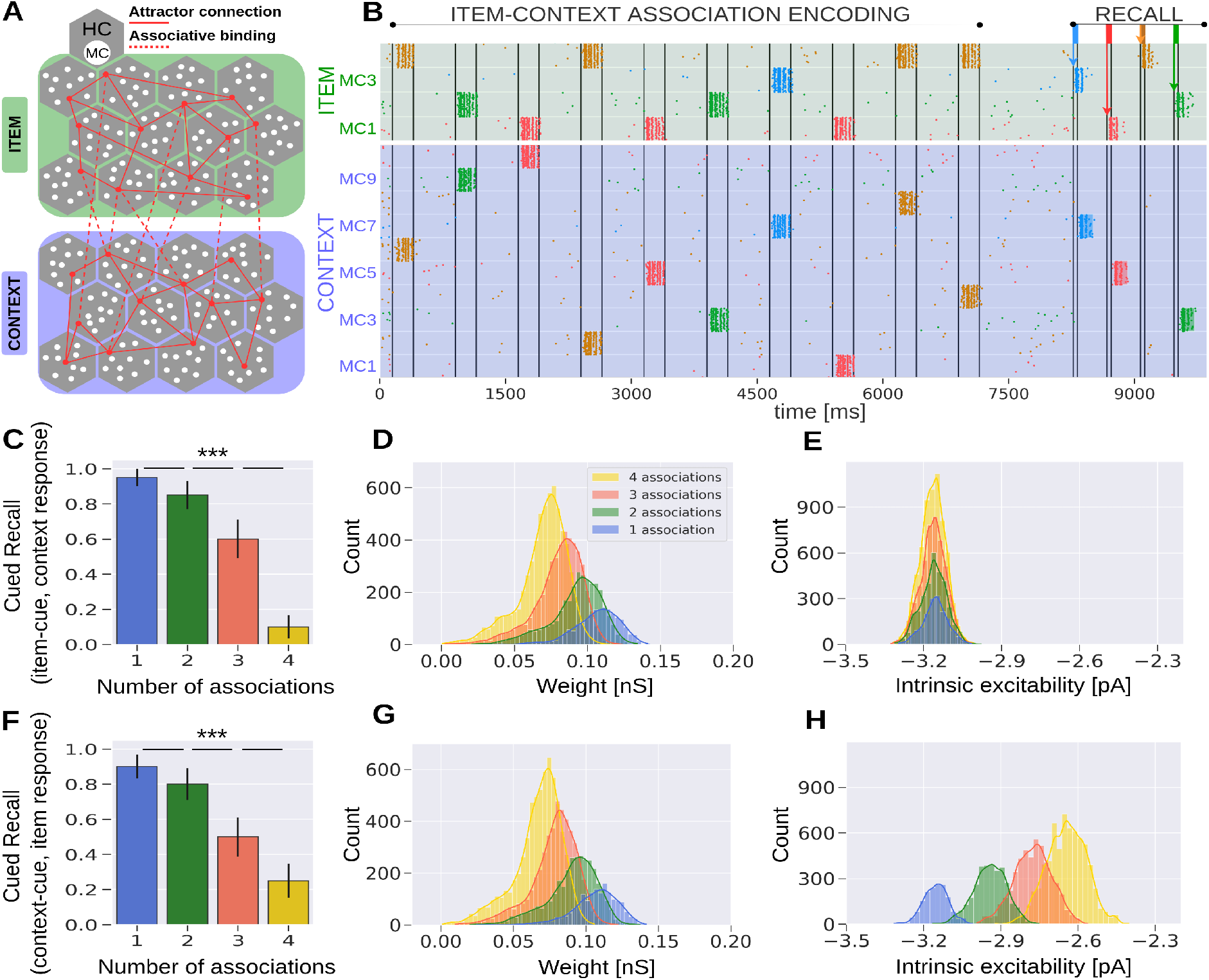
Semantization of episodic memory traces. **A)** Schematic of the Item (green) and Context (blue) networks. Attractor projections are long-range connections across HCs in the same network and learned associative binding are connections between networks. **B)** Spike raster of pyramidal neurons in HC1 of both the Item and Context networks. Items and their corresponding context representations are simultaneously cued in their respective networks (cf. Fig. 2A). Each item is drawn with a unique color, while contexts inherit their coactivated item’s color in the raster (i.e., the yellow pattern in the Item network is repeated over four different contexts, forming four separate associations marked with the same color). The testing phase occurs 1 s after the encoding. Brief 50 ms cues of already studied items trigger their activation. Following item activation, we detect evoked attractor activation in the Context network. **C)** Average cued recall performance in the Context network (20 trials). The bar diagram reveals progressive loss of episodic context information (i.e., semantization) over the number of context associations made by individual cued items (cf. Fig. 2A). **D)** Distribution of plastic connection weights between the Item and Context networks (NMDA component shown here). Weights are noticeably weaker for items which participate in multiple associations. The distributions of synaptic weights exhibit a broader range for the items with multiple context associations, as the sample size is larger. **E)** The distribution of intrinsic excitability currents of pyramidal cells coding for specific context representations. The intrinsic excitability distributions feature similar means because each context is activated exactly once, regardless of whether the associated item forms multiple associations or not. **F)** Average cued recall performance in the Item network (20 trials). Decontextualization over the number of associations is also observed when we briefly cue episodic contexts instead (cf. Fig. 2B, S3). **G)** Distribution of strength of plastic connections from the contexts to their associated items. Analogously to **D**), synapses weaken once an item is encoded in another context. **H)** Intrinsic plasticity distribution of cells in the Item network. Means of the intrinsic excitability distributions are higher for pyramidal cells coding for repeatedly activated items. ***p<0.001 (Mann-Whitney, *N*=20 in **C, F**); Error bars in **C, F** represent standard deviations of Bernoulli distributions; Means of distributions of one, two, three, and four associations in **D, G, H** show significant statistical difference (p<0.001, Mann-Whitney, *N*=2000).

Our simulation results are in line with related behavioral studies (Opitz, 2010; Smith and Manzano, 2010; Smith and Handy, 2014), which also reported item-context decoupling as the items were presented across multiple contexts. In agreement with our study, Opitz (2010) concluded that repetition of an item across different contexts (similar to high context variability) leads to item-context decoupling. Furthermore, Smith and Manzano (2010) demonstrated in an episodic context variability task configuration, that episodicity deteriorates with context overload (number of words per context). Mean recall drops from ∼0.65 (one word per context) to 0.50 (three words per context), reaching ∼0.33 in the most overloaded scenario (fifteen words per context).

In Figure 3E we show the distribution of intrinsic excitability over units representing different contexts. Pyramidal neurons in the Context network have a similar intrinsic excitability, regardless of their selectivity because all the various contexts are encoded exactly once.

Next, analogously to the previous analysis, we show that item-context decoupling emerges also when we briefly cue contexts rather than items during recall testing (Fig. 2B, Fig. S3). In agreement with experimental data (Smith and Manzano, 2010; Smith and Handy, 2014) we obtain evidence of semantization as items learned across several discrete contexts are hardly retrieved when one of their associated contexts serves as a cue (Fig. 3F). We further sample and present the underlying associative weight distribution, between the Context and the Item networks (Fig. 3G). The distributions again reflect the semantization effect in a significant weakening of the corresponding weights. In other words, an assembly of pyramidal neurons representing items encoded across multiple contexts receive weaker projections from the Context network. Beyond four or more associations, the item-context binding becomes so weak that it fails to deliver sufficient excitatory current to trigger associated representations in the Item network. At the same time, intrinsic excitability of item neurons increases with the number of associated contexts corresponding to how much these neurons were active during the encoding phase [Fig. 3H; cf. Egorov et al. (2002), Tully et al. (2014)].

### 2.2 Item-context interactions under STDP

In this section, we contrast the results obtained with the BCPNN synaptic learning rule with those deriving from the more commonly used STDP learning rule in the same episodic memory task (Fig. 2, see STAR⋆METHODS). The modular network architecture as well as neural properties and embedded memory patterns remain identical, but associative projections between networks are now implemented using a standard STDP synaptic learning rule (Morrison et al., 2008). The parameters of the STDP model are summarized in Table S3.

Figure 4A shows an exemplary spike raster of pyramidal cells in HC1 of both the Item and the Context networks, based on the first variant of the episodic memory task described in Figure 2A. As earlier, items are encoded in a single or in multiple different contexts and they are briefly cued later during recall. A successful item activation may lead to a corresponding activation of its associated information in the Context network. We detect these activations as before (see STAR⋆METHODS), and report the cue-based recall score over the number of associations (Fig. 4B).

**Fig. 4.**
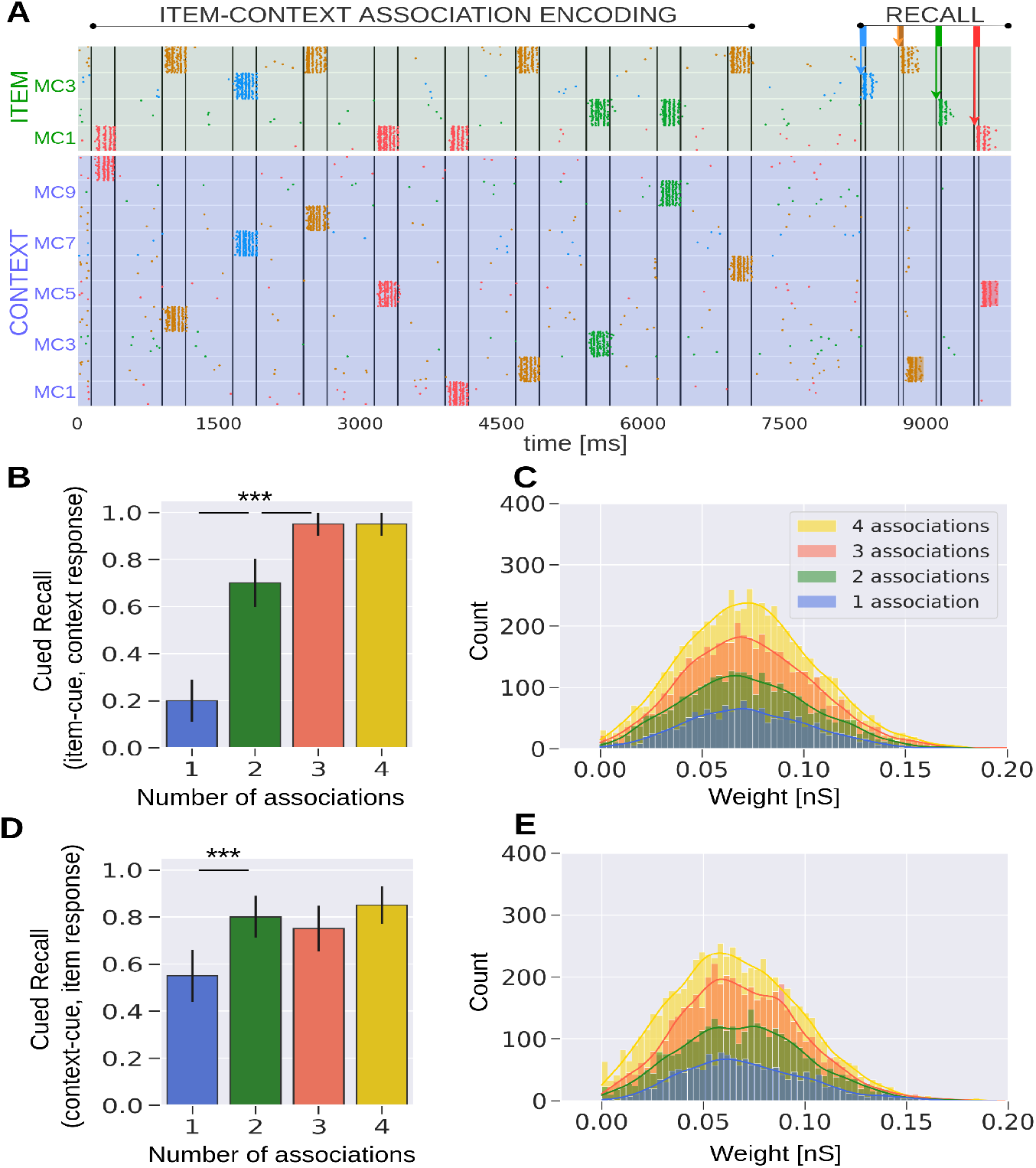
Network model where associative projections are implemented using standard STDP synaptic plasticity. **A)** Spike raster of pyramidal neurons in HC1 of both the Item and Context networks. **B)** Average item-cued recall performance in the Context network (20 trials). Episodic context retrieval is preserved even for high context variability (as opposed to BCPNN, cf. Fig. 3C). **C)** Distribution of NMDA receptor mediated synaptic weights between the item and context neural assemblies following associative binding. The distributions of item-context weights have comparable means at ∼0.065 nS regardless of how many context associations a given item forms. Bins merely display a higher count for the four-association case as the total count of associative weights is more extensive compared to items with fewer associations. **D)** Average cued recall performance in the Item network when episodic contexts are cued (20 trials). **E)** Distribution of NMDA component weights between associated context and item assemblies. ***p<0.001 (Mann-Whitney, *N*=20 in **B, D**); Error bars in **B, D** represent standard deviations of Bernoulli distributions.

Unlike the BCPNN network, we observe no evidence of semantization for high context variability. Instead, recollection is noticeably enhanced with an increasing number of associations, which is in fact the opposite of what would be needed to explain item-context decoupling. STDP generates similarly strong associative binding regardless of context variability (Fig. 4C). The enhanced recollection in high context variability cases stems from the multiplicative effect of synaptic augmentation in the Tsodyks-Makram model on the Hebbian attractor weights. Items stimulated multiple times (e.g., four times) have a higher likelihood of being encoded near the end of the task, leading to more remaining augmentation during testing, thus, effectively boosting cued recall (see Fig. S4). This effect of the enhanced recall diminishes after removing synaptic augmentation from the model (Fig. S5). As far as the context-cued variant of the task is concerned, there are also no signs of item-context decoupling for high context variability (Fig. 4D). The associative projections between Context and Item networks again have distributions with comparable means over context variability (Fig. 4E). An inclusion of intrinsic plasticity dynamics in the model does not explain decontextualization either (see Fig. S6). Overall, decontextualization is not evident in either variant of the episodic memory task under the STDP learning rule.

### 2.3 BCPNN and STDP learning rule in a microcircuit model

To better elucidate the emergent synaptic changes of the BCPNN and STDP model, we also apply these learning rules in a highly reduced microcircuit of spiking neurons. To this end, we now track the synaptic weight changes continuously.

First, we apply the BCPNN learning rule to the microcircuit model. We consider two separate item neurons (ID=1 and 2), which form two or three associations with context neurons (ID=3,4, or 5,6,7), respectively (Fig. 5A). We display the synaptic strength development of the synapse between item neuron-1 and context neuron-3 (two associations, green), as well as the synapse between item neuron-2 and context neuron-5 (three associations, red) over the course of training these associations via targeted stimulation. BCPNN synapses get strengthened when the item-context pairs are simultaneously active and weaken when the item in question is activated with another context. Therefore, synapses of the item neuron that is encoded in three different contexts converge on weaker weights (Fig. 5A, 12 s), than those of the item neuron with two associated contexts. Weight modifications in the microcircuit model reflect the synaptic alterations observed in the large-scale network. BCPNN weights are shaped by traces of activation and co-activation (Eq. 7,8, STAR⋆METHODS), which also get updated during the activation of an item within another context. For example, the item neuron-1 and context neuron-3 are not stimulated together between 6 s and 8 s, but neuron-1 and context neuron-4 are. Thus, the traces of the item activation (*P*_*i*_) increase, while the ones linked to context-3 (*P*_*j*_) decay with a time constant of 15 s (Table S1). Since the item and context neuron (ID=1, 3) are not stimulated together, their coactivation traces (*P*_*i j*_) decay between 6 s and 8 s. Overall, this leads to a weakening of the weight and hence, to a gradual decoupling (Eq. 8, STAR⋆METHODS).

**Fig. 5.**
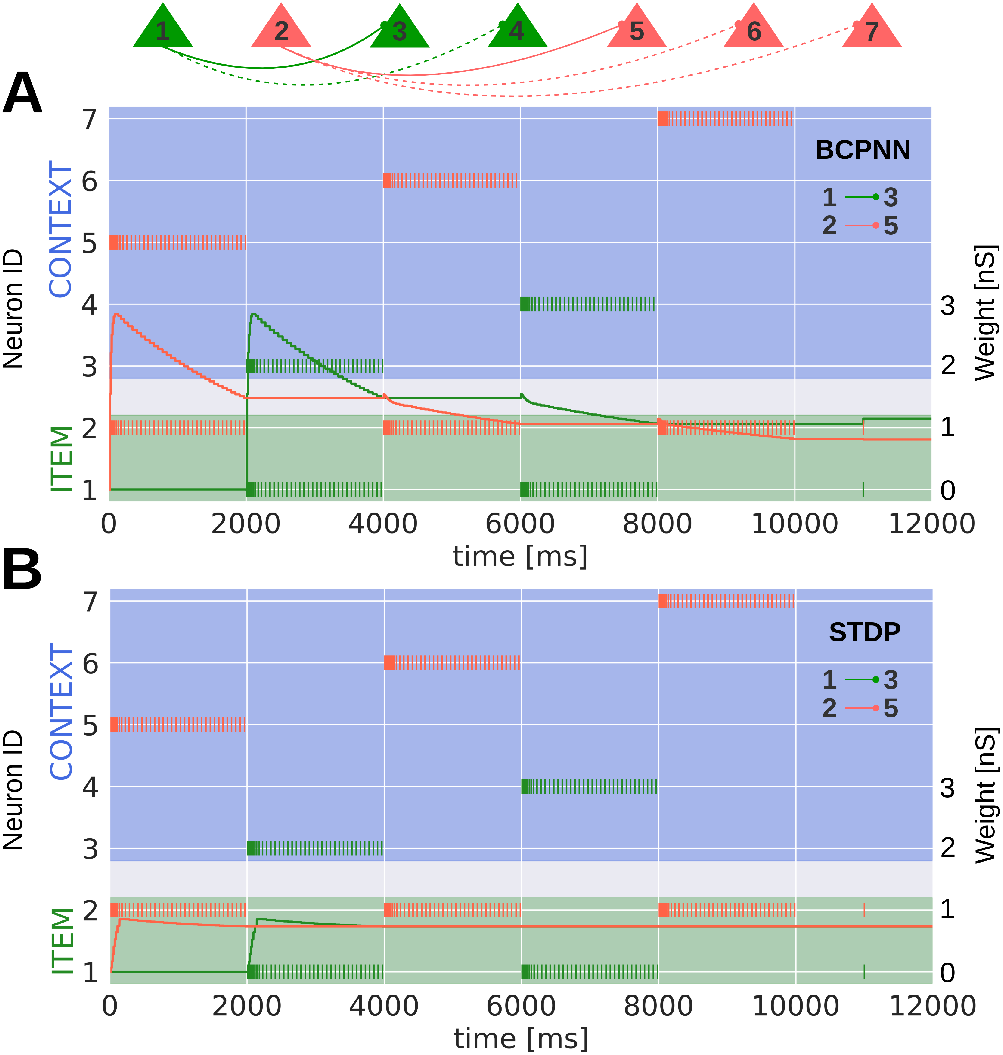
Continuous weight recordings in a microcircuit model with plastic synapses under the **A)** BCPNN or **B)** STDP learning rule. Neural and synaptic parameters correspond to those in the scaled model. In both cases, two item neurons (ID=1,2) are trained to form two or three associations, respectively (dashed connections are simulated but their weight development is not shown here). During training, neurons are stimulated to fire at 20 Hz for 2 s. We display the developing synaptic weight between specific item-context pairs, (ID=1 and 3 in the two-association scenario) and (ID=2 and 5 in the three-association scenario), and compare the converged weight values between the two- and three-association case under both learning rules, following a final readout spike at 11 s.

In the same manner, we keep track of weight change in a microcircuit with the STDP learning rule (Fig. 5B). Unlike the microcircuit with BCPNN presented in Figure 5A, the STDP weights corresponding to the associations made by both item neurons converge to similar values, even though they are associated with different number of contexts. As before, the synapse between an item neuron and an associated context neuron strengthens when this pair is simultaneously active, but remains stable when the item neuron is encoded in another context. For instance, the synapse between item neuron-2 and context neuron-5 strengthens when this pair is encoded (0-2 s), yet remains unaffected when item neuron-2 is activated in another context (i.e., context neuron-6, 4-6 s). This synaptic behavior explains the observed differences between the BCPNN and STDP large-scale model.

### 2.4 Preferential retention

Several studies propose that one-shot salient events promote learning, and that these memories can be retained on multiple time scales ranging from seconds to years (Petrican et al., 2010; Gruber et al., 2016; Frankland et al., 2004; Panoz-Brown et al., 2016; Eichenbaum, 2017; Sun et al., 2018). Hypothetical mechanisms behind these effects are dopamine release and activation of DR1 like receptors, resulting in synapse-specific enhancement (Otmakhova and Lisman, 1996; Kuo et al., 2008), and systems consolidation (McClelland et al., 1995; Fiebig and Lansner, 2014). On the whole, salient or reward driven events may be encoded more strongly as the result of a transient plasticity modulation. Recall from longterm memory is often viewed as a competitive process in which a memory retrieval does not depend only on its own synaptic strength but also on the strength of other components (Shiffrin, 1970). In view of this, we study the effects of plasticity modulation on encoding specific items within particular contexts, with the aim of investigating the role of enhanced learning for semantization in our model.

Using the same network and episodic memory task as before (Fig. 2A), we modulate plasticity during the encoding of item-1 (red) in context-E via *κ*=*κ*_*boost*_ (Eq. 7, STAR⋆METHODS, Table S1). This results in an increased cued recall probability for the item associated with three episodic contexts relative to the unmodulated control (Fig. 6A, Normal vs Biased scenario, 3 associations). Episodic retrieval improves from 0.6 (Normal, Fig. 6A, left) to 0.8 (Biased; modulated plasticity, Fig. 6A, right) when item-1 is cued, which now performs more similarly to item with just two associated contexts. We further analyze and compare the recall of each context when its associated item-1 is cued (Fig. 6B, 3 associations). The control scenario (Normal, Fig. 6B, left) without transient plasticity modulation shows that the three contexts (ID=A, E and J) are all recalled with similar probabilities (20 trials). In contrast, encoding a specific pair with enhanced learning (upregulated *κ*=*κ*_*boost*_) yields higher recall for the corresponding context. In particular, the plasticity enhancement during associative encoding of the context-E (with item-1) results in an increased recall score to 0.8 (0.25 control), while the other associated contexts, ID=A and J, are suppressed (Fig. 6B), primarily due to soft winner-take-all competition between contexts (Fig. 1A).

**Fig. 6.**
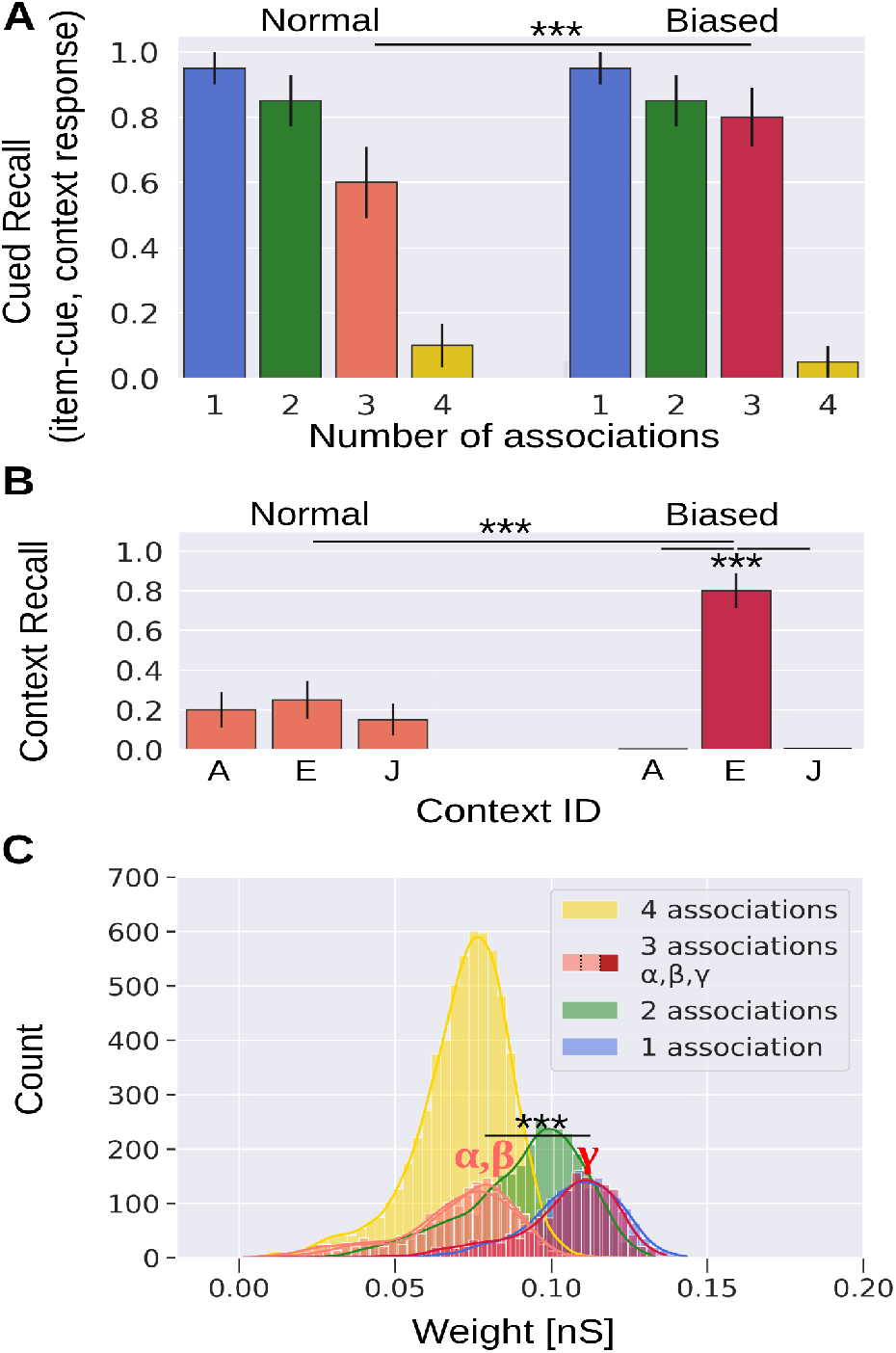
Plasticity modulation of a specific item-context pair enhances recollection and counteracts semantization. **A)** Context recall performance. One of the pairs (context-E, item-1) presented in the episodic memory task (cf. Fig. 2A) is subjected to enhanced plasticity during encoding, resulting in the boosted recall rate (3 associations, Normal vs Biased). **B)** Individual context retrieval contribution in the overall recall (3 associations). Retrieval is similar among the three contexts since plasticity modulation is balanced (left: Normal, *κ*=*κ*_*normal*_, cf. Table S1). However, when context-E is encoded with enhanced learning (with item-1), its recall increases significantly (right: Biased, *κ*=*κ*_*boost*_, cf. Table S1). **C)** Weight distributions of the NMDA weight component. Encoding item-1 with context-E under modulated plasticity yields stronger synaptic weights [3 association, *α,β* (light red, highly overlapping distributions) vs *γ* (dark red)]. ***p<0.001 (Mann-Whitney, *N*=20 in **A, B**, *N*=2000 in **C**); Error bars in **A**,**B** represent standard deviations of Bernoulli distributions; Means of the weight distributions of one, two, three-*α*,-*β*, and four associations in **C** show significant statistical difference (p<0.001, Mann-Whitney, *N*=2000).

We attribute these changes to the stronger weights due to enhanced learning (Fig. 6c, dark red distribution, *γ*). Weights between unmodulated item-context pairs (item-1 and context-A,-J) show mostly unaltered weight distributions (*α,β*, light red), while the biased associative weight distribution between item-1 and context-E is now comparable to the weight distribution of the one-association case. Performance does not exactly match that case though due to some remaining competition among the three contexts. Overall, these results demonstrate how a single salient episode may strengthen memory traces and thus impart resistance to semantization (Rodríguez et al., 2016).

## 3 DISCUSSION

The primary objective of this work was to explore the interaction between synaptic plasticity and context variability in the semantization process. To cast new light on the episodic-semantic interplay, we built a memory model of two spiking neural networks coupled with plastic connections, which collectively represent distributed cortical episodic memory. Our results suggest that some forms of plasticity offer a synaptic explanation for the cognitive phenomenon of semantization, thus bridging scales and linking network connectivity and dynamics with behavior. In particular, we demonstrated that with Bayesian-Hebbian (BCPNN) synaptic plasticity, but not with standard Hebbian STDP, the model can reproduce traces of semantization in the learning outcomes. Notably, this was achieved with biologically constrained network connectivity, postsynaptic potential amplitudes, firing rates and oscillatory dynamics compatible with mesoscale recordings from cortex and earlier models. Nevertheless, our hypothesis of the episodic-semantic interplay at a neural level requires further experimental study of synaptic strength dynamics in particular. We also demonstrated how a transient plasticity modulation (reflecting known isolation effects) may preserve episodicity, staving off decontextualization.

Our study conforms to related behavioral experiments reporting that high context variability or context overload leads to item-context decoupling (Opitz, 2010; Smith and Manzano, 2010; Smith and Handy, 2014). These studies suggest that context-specific memory traces transform into semantic representations while contextual information is progressively lost. Memory traces remain intact but fail to retrieve their associated context. Semantization is typically described as a decontextualization process that occurs over time. However, several experiments, including this study, proposed that exposures of stimuli in different additional contexts (rather than time itself) is the key mechanism advancing semantization (Opitz, 2010; Smith and Manzano, 2010; Smith and Handy, 2014). Admittedly, our hypothesis cannot exclude other seemingly coexisting phenomena that may benefit semantization over time (e.g., reconsolidation or systems consolidation due to sleep or aging).

To our knowledge, there is no other spiking biophysical computational model of comparable detail that captures the semantization of episodic memory explored here, whilst simultaneously offering a neurobiological explanation of this phenomenon. Unlike other dual-process episodic memory models, which require repeated stimulus exposures to support recognition (Norman and O’Reilly, 2003), our model is able to successfully recall events learned in “one shot” (a distinctive hallmark of episodicity). We note that the attractor-based theory proposed in this study does not exclude the possibility of a dual-process explanation for recollection and familiarity (Yonelinas, 2002; Yonelinas et al., 2010).

### 3.1 Related models of familiarity and recollection

Perceptual or abstract single-trace dual-process computational models based on signal detection theory explain episodic retrieval but the potential loss of contextual information is only implied as it does not have its own independent representation (Greve et al., 2010; Wixted, 2007). These computational models often aim to explain traditional R/K behavioral studies. As discussed earlier, participants in such studies are instructed to give a Know response if the stimulus presented in the test phase is known or familiar without any contextual detail about its previous occurrence. Conversely, Remember judgments are to be provided if the stimulus is recognized along with some recollection of specific contextual information pertaining to the study episode. This results in a strict criterion for recollection, as it is possible for a subject to successfully recall an item but fail to retrieve the source information (Ryals et al., 2013). Numerous studies suggest that recollection contaminates Know reports because recalling source information sensibly assumes prior item recognition (Wais et al., 2008; Johnson et al., 2009). Mandler (1979, 1980), and Atkinson and Juola (1973) treat familiarity as an activation of preexisting memory representations. Our results are compatible with this notion because our model proposes to treat item-only activations as Know judgments, while those accompanied by the activation of context representations best correspond to a Remember judgment. Item activation is a faster process and precedes context retrieval (Yonelinas and Jacoby, 1994), and our model reflects this finding by necessity, as item activations are causal to context retrieval.

To us, familiarity recognition is simply characterized by a lack of contextual information, yet the distinction we make between Context and Item networks is arbitrary. Any item can be a context and vice versa, so the networks are interchangeable. While sparse interconnection is sufficient for our model’s function, both networks may just as well be part of the same modality and cortical brain areas. A more specific scenario might assume that items and contexts share part of the same local network. In principle, our model should be capable of replicating similar results in a single modality scenario.

### 3.2 Semantization on longer time scales

Source recall is likely supported by multiple independent, parallel, interacting neural structures and processes since various parts of the medial temporal lobes, prefrontal cortex and parts of the parietal cortex all contribute to episodic memory retrieval including information about both where and when an event occurred (Diana et al., 2007; Gilboa, 2004; Watrous et al., 2013). A related classic idea on semantization is the view that it is in fact an emergent outcome of systems consolidation. Sleep-dependent consolidation in particular has been linked to advancing semantization of memories and the extraction of gist information. (Friedrich et al., 2015; Payne et al., 2009).

Models of long-term consolidation suggest that richly contextualized memories, become more generic over time. Without excluding this possibility, we note that this is not always the case, as highly salient memories often retain contextual information (which our model speaks to). Instead, our model argues for a much more immediate neural and synaptic contribution to semantization that does not require slow multi-area systems level processes that have yet to be specified in sufficient detail to be tested in neural simulations. We have previously shown, however, that an abstract simulation network of networks with broader distributions of learning time constants can consolidate memories across several orders of magnitude in time, using the same Bayesian-Hebbian learning rule as used here (Fiebig and Lansner, 2014). That model included representations for prefrontal cortex, hippocampus, and wider neocortex, implementing an extended complementary learning systems theory (McClelland et al., 1995), which is itself an advancement of systems consolidation (Squire and Alvarez, 1995). We consequently expect that the principled mechanism of semantization explored here can be scaled along the temporal axis to account for lifelong memory, provided that the plasticity involved is itself Bayesian-Hebbian. Our model does not advance any specific anatomical argument as to the location of the respective networks (Diana et al., 2007; Yonelinas, 2002).

The model purposefully relies on a generic cortical architecture focused on a class of synaptic plasticity mechanisms which may well serve as a substrate of a wider system across brain areas and time.

### 3.3 Biological plausibility and parameter sensitivity

We investigate and explain behavior and macroscale system dynamics with respect to neural processes, biological parameters of network connectivity, and electrophysiological evidence. Our model consequently builds on a broad range of biological constraints such as intrinsic neuronal parameters, cortical laminar cell densities, plausible delay distributions, and network connectivity. The model reproduces plausible postsynaptic potentials (EPSPs, IPSPs) and abides by estimates of connection densities (i.e., in the associative pathways and projections within each patch), axonal conductance speeds, typically accepted synaptic time constants for the various receptor types (AMPA, NMDA, and GABA), with commonly used neural and synaptic plasticity time constants (i.e., adaptation, depression). We reproduce oscillatory dynamics in multiple frequency ranges, that were previously studied in the same modular spiking network implementations (Lundqvist et al., 2010, 2011; Herman et al., 2013).

The model synthesizes a number of functionally relevant processes, embedding different components to model composite dynamics, hence, it is beyond this study to perform a detailed sensitivity analysis for every parameter. Instead, we provide insightful observations for previously unexplored parameters that may critically affect semantization. Importantly, a highly related modular cortical model already investigated sensitivity to important short-term plasticity parameters (Fiebig and Lansner, 2017). After extensive simulation testing, we conclude that the model is generally robust to a broad range of parameter changes and degrades gracefully. Small networks like this are typically more sensitive to parameter changes, so conversely, we expect even lower sensitivity to parameter variations in a full scale system.

The P trace decay time constant, *τ*_*p*_, of the BCPNN model is critical for the learning dynamics modelled in this study because it controls the speed of learning in associative connections. High values of *τ*_*p*_ lead to slower and more long-lasting learning. Varying *τ*_*p*_ by *±*30% does not change the main outcome, that is, episodicity still deteriorates with a higher context variability. Slower weight development may result in weaker associative binding and overall lower recall though (and vice versa for faster learning). To compensate for this loss of episodicity, an additional increase in the unspecific input is usually sufficient to trigger comparable recall rates. Alternatively, the recurrent excitatory gain can be amplified to complete noisy inputs towards discrete embedded attractors. Unspecific background input during recall plays a critical role as well. We use a low such noise input to model cue-association responses, however, when boosted by +40%, the model operates in a free replay regime instead, where cues become unnecessary as the network retrieves content without them by means of intrinsic background noise.

This study also demonstrated how a selective transient increase of plasticity can counteract semantization. The plasticity of the model can be modulated via the parameter *κ* (Eq. 7, STAR⋆METHODS). Typically, *κ* is set to 1 (*κ*=*κ*_*normal*_), whereas we double plasticity (*κ*=*κ*_*boost*_), when modelling salient episodic encoding. We noticed that by selectively tripling or quadrupling plasticity (relative to baseline) during encoding of a specific pair whose item component forms many other associations, the source recall improves progressively (data shown only for *κ*=*κ*_*boost*_ in Sect. 2.4).

Finally, in Section 2.3 we compared STDP and BCPNN plasticity in a highly reduced model. We bind items with contexts to form different number of associations and keep track of the weight development per time step. STDP plasticity generated same magnitude item-context binding regardless of how many associations an item forms. A detailed parameter analysis for every critical synaptic parameter (*±*30%) did not yield any behaviorally significant changes to the converged weights.

### 3.4 Conclusions

We have presented a computational mesoscopic network model to examine the interplay between episodic and semantic memory with the grand objective to explain mechanistically the semantization of episodic traces. Compared to other models of episodic memory, which are typically abstract, our model, built on various biological constraints (i.e., plausible postsynaptic potentials, firing rates, etc.) accounting for neural processes and synaptic mechanisms, emphasizes the role of synaptic plasticity in episodic forgetting. Hence it bridges micro and mesoscale mechanisms with macroscale behavior and dynamics. In contrast to standard Hebbian learning, our Bayesian version of Hebbian learning readily reproduced prominent traces of semantization.

## Acknowledgments

This research was supported by Vetenskapsrådet 2018-05360 and 2016-05871, the Swedish e-Science Research Centre (SeRC), Digital Futures, and European Commission H2020 program. The simulations were performed on resources provided by Swedish National Infrastructure for Computing (SNIC) at the PDC Center for High Performance Computing, KTH Royal Institute of Technology.

## STAR⋆METHODS

Detailed methods are provided in the online version of this paper and include the following:

- KEY RESOURCES TABLE
- METHODS DETAILS
  - Spike-based BCPNN plasticity
  - Spike-based STDP learning rule
  - Two-network architecture and connectivity
  - Axonal conduction delays
  - Stimulation Protocol
  - Attractor activation detector
  - Simulation Tools

## 4 STAR⋆METHODS

### 4.1 KEY RESOURCES TABLE

**Table 1.**
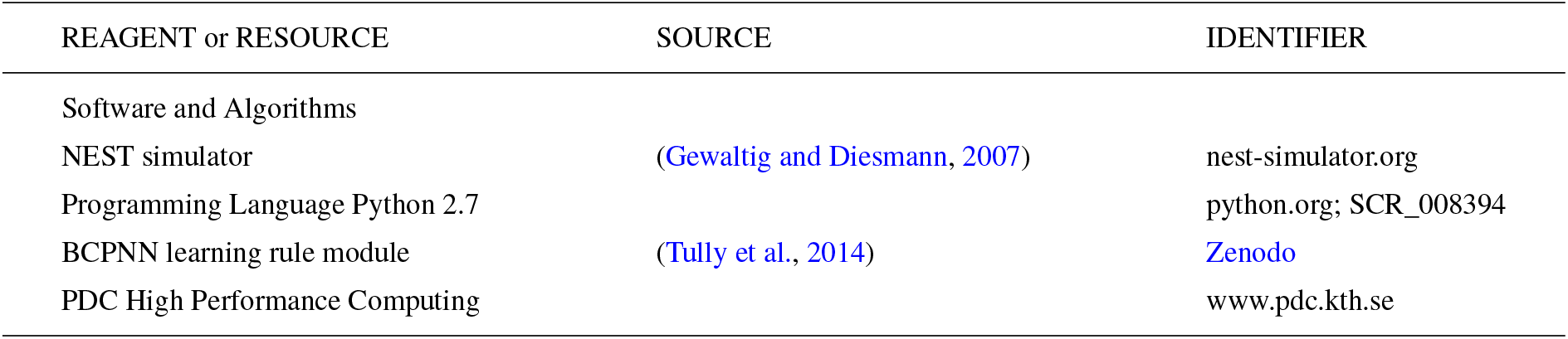

### 4.2 METHODS DETAILS

#### 4.2.1 Neuron and synapse model

We use adaptive exponential integrate-and-fire point model neurons, which feature spike frequency adaptation, enriching neural dynamics and spike patterns, especially for the pyramidal cells (Brette and Gerstner, 2005). The neuron model offers a good phenomenological description of typical neural firing behavior, but it is limited in predicting the precise time course of the sub-threshold membrane voltage during and after a spike or the underlying biophysical causes of electrical activity (Gerstner and Naud, 2009). We slightly modified it for compatibility with the BCPNN synapse model (Tully et al., 2014) by integrating an intrinsic excitability current.

Development of the membrane potential *V*_*m*_ and the adaptation current *I*_*w*_ is described by the following equations:

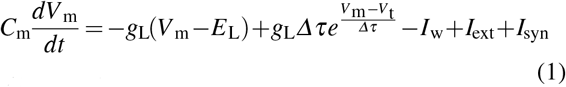

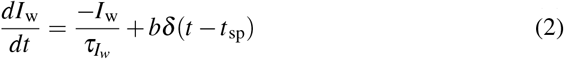

Equation 1 describes the dynamics of the membrane potential *V*_*m*_ including an exponential voltage dependent activation term. A leak current is driven by the leak reversal potential *E*_*L*_ through the conductance *g*_*L*_ over the neural surface with a capacity *C*_*m*_. Additionally, *V*_*t*_ is the spiking threshold, and *Δ*_T_ shapes the spike slope factor. After spike generation, membrane potential is reset to *V*_*r*_. Spike emission upregulates the adaptation current by *b*, which recovers with time constant *τ*_*Iw*_ (Table S1). We neglect subthreshold adaptation, which is part of some AdEx models.

Besides a specific external input current *I*_*ext*_, model neurons receive synaptic currents 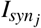 from conductance based glutamatergic and GABA-ergic synapses. Glutamatergic synapses feature both AMPA/NMDA receptor gated channels with fast and slow conductance decay dynamic, respectively. Current contributions for synapses are described as follows:

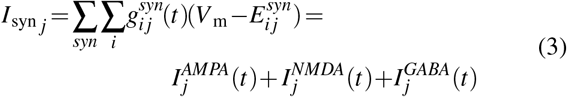

The glutamatergic synapses are also subject to synaptic depression and augmentation with a decay factor *τ*_*D*_ and *τ*_*A*_, respectively (Table S1), following the Tsodyks-Markram formalism (Tsodyks and Markram, 1997). The utilization factor *U*, encodes variations in the release probability of available resources:

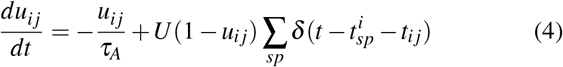

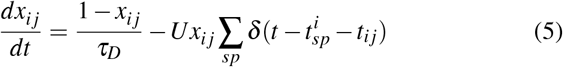

#### 4.2.2 Spike-based BCPNN plasticity

We implement synaptic plasticity of AMPA and NMDA connection components using the BCPNN learning rule (Lansner and Ekeberg, 1989; Wahlgren and Lansner, 2001; Tully et al., 2014). BCPNN is derived from Bayes rule, assuming a postsynaptic neuron employs some form of probabilistic inference to decide whether to emit a spike or not. In general, it is considered more complex than the standard STDP learning rule (Caporale and Dan, 2008), and it reproduces the main features of STDP plasticity. As other spiking synaptic learning rules, it is so far insufficiently validated against quantitative experimental data on biological synaptic plasticity.

The BCPNN synapse continuously updates three synaptic biophysically plausible local memory traces, *P*_*i*_, *P*_*j*_ and *P*_*i j*_, implemented as exponentially moving averages (EMAs) of pre-, post- and co-activation, from which the Bayesian bias and weights are calculated. EMAs prioritize recent patterns, so that newly learned patterns gradually replace old memories. Specifically, learning implements a three-level procedure of exponential filters which defines Z, E and P traces. E traces, which enable delayed reward learning, are not used here because such conditions are not applicable to the modelled task.

To begin with, BCPNN receives a binary sequence of pre- and postsynaptic spiking events (*S*_*i*_, *S* _*j*_) to calculate the traces *Z*_*i*_ and *Z*_*j*_:

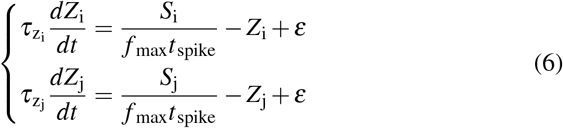

*f*_*max*_ denotes the maximal neuronal spike rate, *ε* is the lowest attainable probability estimate, *t*_*spike*_ denotes the spike duration while 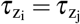 are the pre and postsynaptic time constants, respectively (5 ms for AMPA, and 100 ms for NMDA components, Table S1).

P traces are then estimated from the Z traces as follows:

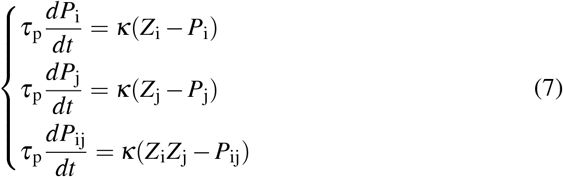

The parameter *κ* adjusts the learning rate, reflecting the action of endogenous modulators of learning efficacy (i.e., activation of a D1R-like receptor). Setting *κ*=0 freezes the network’s weights and biases, though in our simulations the learning rate remains constant (*κ*=1) during encoding (Sect. 2.1, 2.2). However, we trigger a transient increase of plasticity in specific scenarios to model preferential retention, assuming encoding of salient events (Sect. 2.4 and Table S1).

Finally, *P*_*i*_, *P*_*j*_ and *P*_*i j*_ are used to calculate intrinsic excitability *β* _*j*_ and synaptic weights *w*_*i j*_ with a scaling factor *β*_*gain*_ and 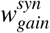 respectively (Table S1):

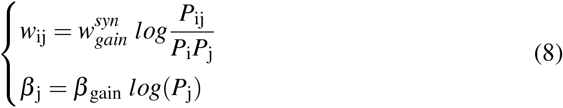

#### 4.2.3 Spike-based STDP learning rule

In our study, we examine the impact on semantization when the STDP learning rule replaces BCPNN associative connectivity in the same episodic memory task. Synapses under STDP are developed and modified by a repeated pairing of pre- and postsynaptic spiking activity, while their relative time window shapes the degree of modification (Ren et al., 2010). The amount of trace modification depends on the temporal difference (*Δ*_*t*_) between the time point of the presynaptic action potential (*t*_*i*_) and the occurrence of the postsynaptic spike (*t*_*j*_) incorporating a corresponding transmission delay from neuron i to j (*τ*_*d*_):

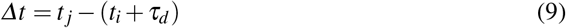

After processing *Δ t*, STDP updates weights accordingly:

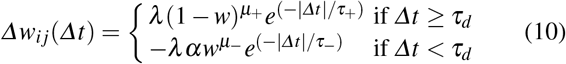

Here, *λ* corresponds to the learning rate, *α* reflects a possible asymmetry between the scale of potentiation and depression, *τ*_*±*_ control the width of the time window, while *µ*_*±*_ ∈ {0,1} allows to choose between different versions of STDP (i.e., additive, multiplicative), (Morrison et al., 2008). Synapses are potentiated if the synaptic event precedes the postsynaptic spike and get depressed if the synaptic event follows the postsynaptic spike (Van Rossum et al., 2000).

Associative weights *w*_*i j*_ are initialized to *w*_0_, and their maximum allowed values are constrained according to *w*_*max*_ to ensure that synaptic weights are always positive and between [*w*_0_, *w*_*max*_] (Table S3). The resulting associative weight distributions are generally comparable in strength to the BCPNN model weights, but to make them match, we adjust *w*_*max*_ in conjunction with a reasonably small learning rate *λ*. To obtain a stable competitive synaptic modification, the integral of *Δ w*_*i j*_ must be negative (Song et al., 2000). To ensure this, we choose *α*=1.2, which introduces an asymmetry between the scale of potentiation and depression along with a symmetric time window resulting in a ratio of *ατ*_*-*_*/τ*_+_>1.0 (Ren et al., 2010). We set *µ*_*±*_=1 resulting in multiplicative STDP (in between values lead to rules which have an intermediate dependence on the synaptic strength). Pyramidal cells receive an unspecific background noise at 420 Hz during recall.

#### 4.2.4 Two-network architecture and connectivity

The network model includes two reciprocally connected networks, the Item and Context networks. For simplicity, we assumed that item and context information engage different modalities and cortical areas and thus the corresponding networks are located at a substantial distance (Table S2). Both networks span a regular-spaced grid of 12 HCs (Table S2), each with a diameter of 500 *µ*m (Mountcastle, 1997). Our model employs distributed orthogonal representations with one active MC per HC, approximating the exceedingly sparse neocortical activity patterns with marginal overlap. Each minicolumn is composed of 30 pyramidal cells with shared selectivity, forming a functional (not strictly anatomical) column. In total, the 24 HCs of the model contain 7200 excitatory and 480 inhibitory cells, significantly downsampling the number of MC per HC (∼100 MC per HC in biological cortex). The high degree of recurrent connectivity within MCs (Thomson et al., 2002; Yoshimura and Callaway, 2005) and between them link coactive MCs into larger cell assemblies (Eyal et al., 2018; Binzegger et al., 2009; Muir et al., 2011; Stettler et al., 2002). Long-range bidirectional inter-area connections (item-context bindings or associative connections) are plastic (shown in Fig. 1A only for MC1 in HC1 of the Context network), binding items and contextual information (Ranganath, 2010). Recurrent connectivity establishes 100 active plastic synapses on average onto each pyramidal cell from other pyramidals with the same selectivity, due to a sparse inter-area connectivity (*cp*_*PPA*_) and denser local connectivity (*cp*_*PP*_, *cp*_*PPL*_; connection probabilities are indicated in Fig. 1A only for MC1 in HC1 of the Context network). The model yields biologically plausible excitatory postsynaptic potentials (EPSPs) for connections within HCs (0.45 *±* 0.13 mV), measured at resting potential *E*_*L*_ (Thomson et al., 2002). Densely recurrent non-specific monosynaptic feedback inhibition mediated by fast spiking inhibitory cells (Kirkcaldie, 2012) implements a local winner-take-all structure (Binzegger et al., 2009) amongst the functional columns. Inhibitory post-synaptic potentials (IPSPs) have an amplitude of -1.160 mV (*±*0.003) measured at -60 mV (Thomson et al., 2002). These bidirectional connections between basket and pyramidal cells within the local HCs are drawn with a 70% connection probability. Notably, double bouquet cells shown in Figure 1A, are not explicitly simulated, but their effect is nonetheless expressed by the BCPNN rule. A recent study based on the same basic model architecture demonstrated that learned mono-synaptic inhibition between competing attractors is functionally equivalent to the disynaptic inhibition mediated by double bouquet and basket cells (Chrysanthidis et al., 2019). Parameters characterising other neural and synaptic properties including BCPNN can be found in Table S1.

Figure 1B shows the weight distributions of embedded distributed cell assemblies, representing different memories stored in the Item and Context networks. Attractor projections can be further categorized into strong local recurrent connectivity within HCs, and slightly weaker long-range excitatory projections across HCs (Fig. 1C).

#### 4.2.5 Axonal conduction delays

Conduction delays (*t*_*i j*_) between a presynaptic neuron *i* and a postsynaptic neuron *j* are calculated based on their Euclidean distance, *d*, and a conduction velocity *V* (Eq. 11). Delays are randomly drawn from a normal distribution with a mean according to distance and conduction velocity, with a relative standard deviation of 30% of the mean. In addition, a minimal delay of 1.5 ms (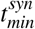, Table S2) is added to reflect synaptic delays due to effects that are not explicitly modelled, e.g. diffusion of neurotransmitters over the synaptic cleft, dendritic branching, thickness of the cortical sheet and the spatial extent of columns. Associative inter-area projections have a tenfold faster conduction speed than those within each network, reflecting axonal myelination.

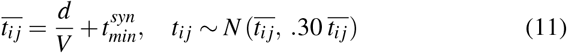

#### 4.2.6 Stimulation Protocol

Noise input to pyramidal cells and fast spiking inhibitory basket cells is generated by two independent Poisson generators with conductances of opposing signs. Pyramidal cells coding for specific items and contexts are stimulated with an additional specific excitation during encoding and cued recall (all parameters in Table S2). Item-context association encoding is preceded by a brief period of background noise excitation to avoid initialization transients.

#### 4.2.7 Attractor activation detector

We detect and report cue-based activation of items or contexts by utilizing an attractor activation detection algorithm based on EMAs of spiking activity. Pattern-wise EMAs are calculated using Equation 12, where the delta function *δ* denotes the spike events of a pattern-selective neural population of *n*_*pop*_=30 pyramidal cells. The filter time constant *τ*=40 ms is much larger than the sampling time interval *Δ T* =1 ms.

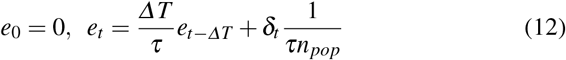

Pattern activations are detected by a simple threshold (*r*_*th*_) at about tenfold the baseline activity with a small caveat: To avoid premature offset detection due to synchrony in fast spiking activity, we only count activations as terminated if they do not cross the threshold again in the next 40 ms. Despite the complications of nested oscillations, this method is highly robust due to the explosive dynamics of recurrent spiking activity for activated attractors in the network. Any attractor activation that crosses this threshold for at least 40 ms is considered a successful recall.

## 5 SUPPLEMENTARY MATERIAL

**Fig. S1.**
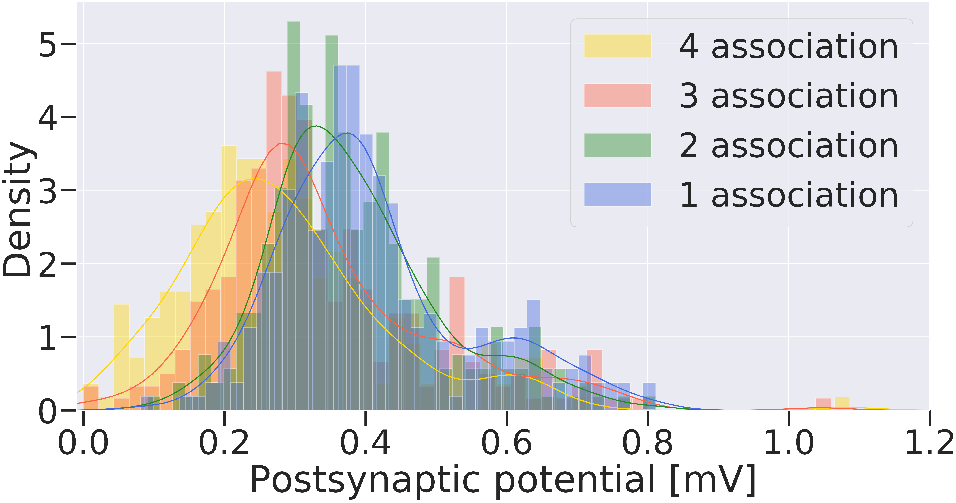
Excitatory postsynaptic potentials (EPSPs) for the binding between Item and Context networks. EPSPs were recorded (at resting potential *E*_*L*_) after item-context association encoding phase. We stimulate individually all the neurons in HC1 of an item which forms one, two, three or four associations and record the postsynaptic potential onto their associated context neurons. Means of the EPSP distributions show significant statistical difference (*p*<0.05 for one vs two associations; *p*<0.001 for two vs three and three vs four associations, Mann-Whitney, *N*=300).

**Fig. S2.**
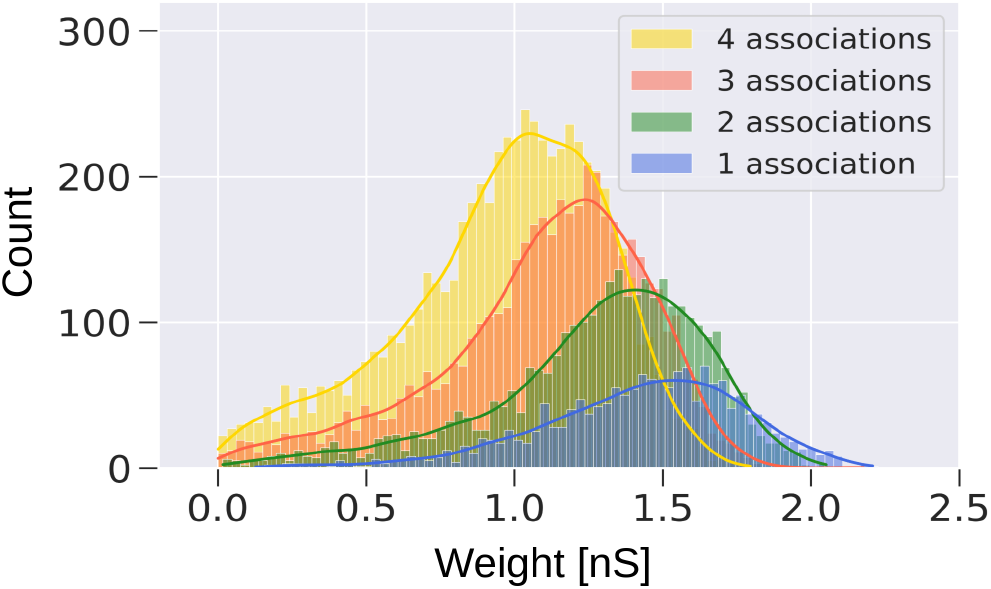
Distributions of the AMPA component weights between Item and Context networks. Slower NMDA receptor weights follow a similar pattern of weakening for items which participate in multiple associations. Means of the weight distributions of one, two, three, and four associations show significant statistical difference (p<0.001, Mann-Whitney, *N*=2000).

**Fig. S3.**
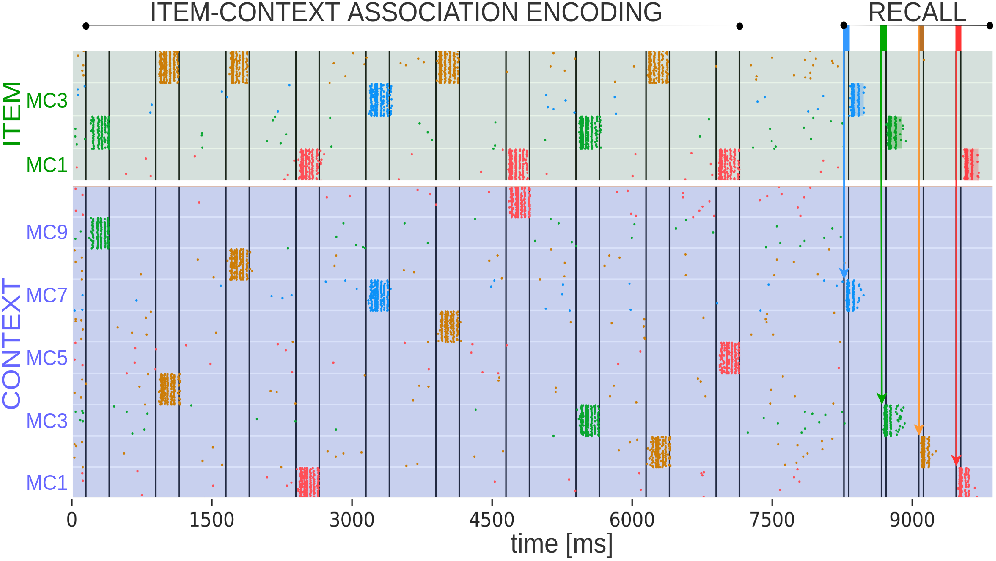
Spike raster of pyramidal cells in HC1 of both the Item and Context networks in the BCPNN model. Items and their corresponding context representations are simultaneously cued in their respective networks. The testing phase occurs 1 s after the encoding and triggers activations via partial cues of contexts (50 ms cues). Repetition of items across various contexts leads to progressive item-context decoupling. Item-4 is repeated across four different contexts, and while its associated context gets activated when cued (context-B), item-4 is not retrieved.

**Fig. S4.**
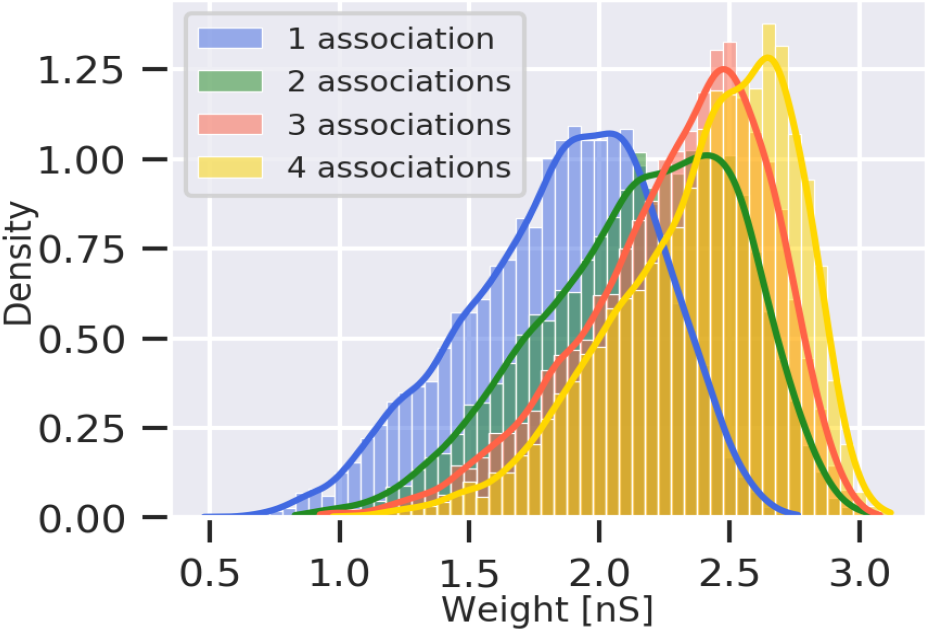
Weight distribution of AMPA component weights of the Item network including synaptic augmentation. The multiplicative effect of synaptic augmentation on the consolidated Items features stronger combined synaptic strength for items with higher context variability. Slower NMDA receptor weights follow a similar pattern. Means of the weight distributions of one, two, three, and four associations show significant statistical difference (p<0.001, Mann-Whitney, *N*=2000).

**Fig. S5.**
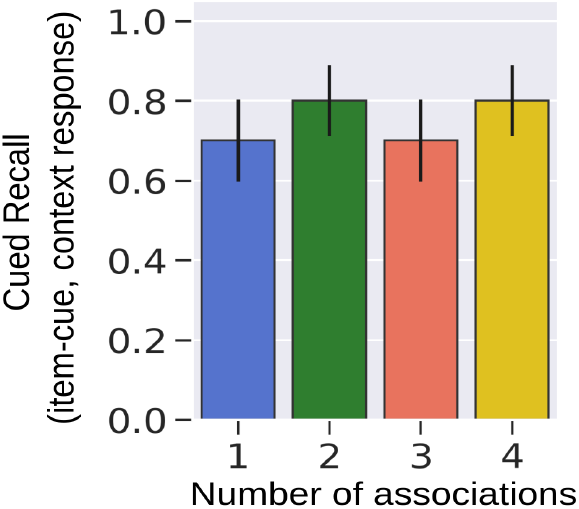
Cued recall under STDP after removing synaptic augmentation. Average item-cued recall performance in the Context network (20 trials). To compensate for the removal of augmentation, we increased the stimulation rates and the synaptic gain leading to comparable elicited spiking activity. Error bars represent standard deviations of Bernoulli distributions.

**Fig. S6.**
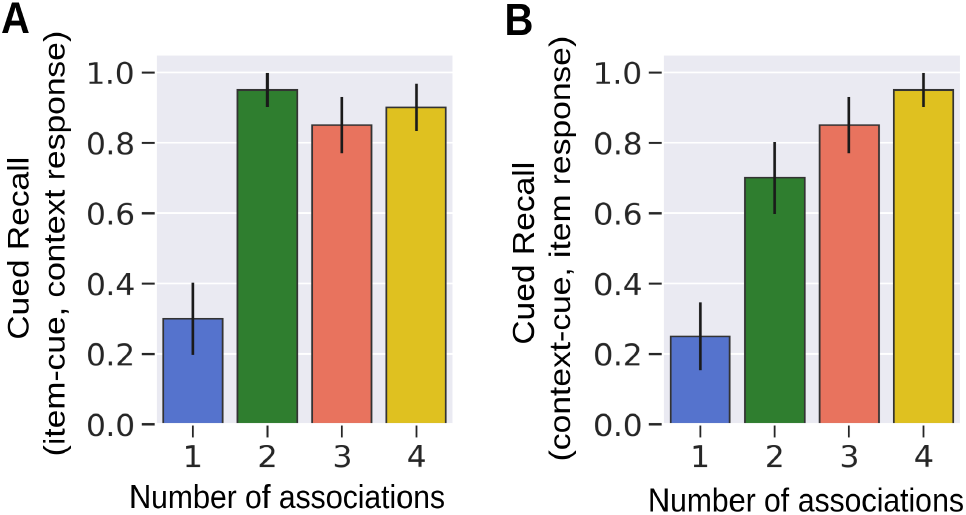
Cued recall under STDP including intrinsic plasticity. **A**) Average item-cued recall performance in the Context network (20 trials). **B**) Average item-cued recall performance in the Context network. Episodic context retrieval is enhanced for high context variability predominantly because of intrinsic excitability dynamics and synaptic augmentation. We observe an opposite trend to the decontextualization effect seen in Figure 3C. Error bars represent standard deviations of Bernoulli distributions.

**Table S1.** Model and BCPNN parameters

| Parameter | Symbol | Value | Parameter | Symbol | Value | Parameter | Symbol | Value |
| --- | --- | --- | --- | --- | --- | --- | --- | --- |
| Adaptation current | $b$ | 86 pA | Utilization factor | $U$ | 0.2 | BCPNN AMPA gain | $w_{gain}^{AMPA}$ | 0.76 nS |
| Adaptation decay time constant | $\tau_w$ | 280 ms | Augmentation decay time constant | $\tau_A$ | 5 s | BCPNN NMDA gain | $w_{gain}^{NMDA}$ | 0.07 nS |
| Membrane capacitance | $C_m$ | 280 pF | Depression decay time constant | $\tau_D$ | 280 ms | BCPNN bias current gain | $\beta_{gain}$ | 40 pA |
| Leak reversal potential | $E_L$ | -70.6 mV | AMPA synaptic time constant | $\tau^{AMPA}$ | 5 ms | BCPNN lowest spiking rate | $f_{min}$ | 0.2 Hz |
| Leak conductance | $g_L$ | 14 nS | NMDA synaptic time constant | $\tau^{NMDA}$ | 100 ms | BCPNN highest spiking rate | $f_{max}$ | 25 Hz |
| Upstroke slope factor | $\Delta_T$ | 3 mV | GABA synaptic time constant | $\tau^{GABA}$ | 5 ms | BCPNN lowest probability | $\epsilon$ | 0.01 |
| Spike threshold | $V_t$ | -55 mV | AMPA reversal potential | $E^{AMPA}$ | 0 mV | BCPNN Spike event duration | $t_{spike}$ | 1 ms |
| Spike reset potential | $V_r$ | -60 mV | NMDA reversal potential | $E^{NMDA}$ | 0 mV | P trace time constant | $\tau_p$ | 15 s |
| Refractory period | $\tau_{ref}$ | 5 ms | GABA reversal potential | $E^{GABA}$ | -75 mV | Modulated plasticity | $\kappa_{boost}$ | 2 |
| | | | Regular plasticity | $\kappa_{normal}$ | 1 | | | |

**Table S2.** Network layout, connectivity and stimulation protocol

| Layout | Symbol | Value | Connectivity | Symbol | Value | Stimulation | Symbol | Value |
| --- | --- | --- | --- | --- | --- | --- | --- | --- |
| Cortical patch size | $C_{ps}$ | 2.0 x 1.5 mm | Axonal Conduction Speed | $V$ | 0.2 m/s | Background noise PYR (encoding) | $r_{bg-encoding}^{PYR}$ | 650 Hz |
| Simulated HCs | $n_{HC}$ | 12 | Myelinated axonal speed | $V_{myel}$ | 2 m/s | Background noise PYR (recall) | $r_{bg-recall}^{PYR}$ | 450 Hz |
| Simulated MCs | $n_{MC}$ | 120 | Minimal synaptic delay | $t_{min}^{syn}$ | 1.5 ms | Background noise BA | $r_{bg}^{BA}$ | 75 Hz |
| Simulated MCs per HC | $n_{MC}^{HC}$ | 10 | Hypercolumn diameter | $d_{HC}$ | 0.5 mm | Background conductance | $g_{bg}^{PYR,BA}$ | $\pm 1.5$ nS |
| No. of items | $n_{ITEM}$ | 4 (from 10) | Distance between networks | $d_{CONTEXT}^{ITEM}$ | 10 mm | Stimulation duration | $t_{stim}$ | 250 ms |
| No. of contexts | $n_{CONTEXT}$ | 10 (from 10) | PYR-PYR recurrent cp | $cp_{PP}$ | 0.2 | Stimulation rate | $r_{stim}$ | 500 Hz |
| Layer 2/3 pyramidal per MC | $n_{MC}^{PYR-L23}$ | 30 | PYR-PYR long-range cp | $cp_{PPL}$ | 0.25 | Cue stimulation length | $t_{cue}$ | 50 ms |
| Basket cells per MC | $n_{MC}^{Basket}$ | 2 | PYR-PYR associative cp | $cp_{PPA}$ | 0.02 | Cue stimulation rate | $r_{cue}$ | 400 Hz |
| MC grid size (Item + Context) | $G_{MC}^{TOTAL}$ | 24 x 10 | PYR-BA cp, BA-PYR cp | $cp_{PB}, cp_{BA}$ | 0.7 | Stimulation and cue conductance | $g_{stim}$ | +1.5 nS |
| | | | PYR-BA cc | $g_{PB}$ | 3 nS | Interstimulus interval | $T_{stim}$ | 500 ms |
| | | | BA-PYR cc | $g_{BP}$ | -7 nS | Attractor detection threshold | $r_{th}$ | 10 Hz |

PYR, Pyramidal cell; BA, Basket cell. cp, connection probability; cc, connection conductance.

**Table S3.** STDP synaptic model parameters

| Parameter | Symbol | Value |
| --- | --- | --- |
| Weight initialization | $w_0$ | 0 nS |
| AMPA maximum allowed weight | $w_{max}^{AMPA}$ | 13.5 nS |
| NMDA maximum allowed weight | $w_{max}^{NMDA}$ | 3.5 nS |
| Learning rate | $\lambda$ | 0.01 |
| Asymmetry parameter | $\alpha$ | 1.2 |
| Weight dependence exponent, potentiation | $\mu_+$ | 1 |
| Weight dependence exponent, depression | $\mu_-$ | 1 |
| Symmetric time window | $\tau_{\pm}$ | 20 ms |

